# Dissecting the sublexical route for reading: Frontal and parietal networks support learned orthography-to-phonology mappings

**DOI:** 10.1101/2025.02.27.640623

**Authors:** Sara M. Dyslin, Andrew T. DeMarco, Ryan Staples, J. Vivian Dickens, Sarah F. Snider, Rhonda B. Friedman, Peter E. Turkeltaub

**Affiliations:** Department of Neurology, Georgetown University Medical Center, 4000 Reservoir Rd NW, Building D, Washington, DC 20057; Department of Rehabilitation Medicine, Georgetown University Medical Center, 4000 Reservoir Rd NW, Building D, Washington, DC 20057; MedStar National Rehabilitation Hospital, 102 Irving St NW, Washington, DC 20010

**Author notes:** **Corresponding Author:** Sara M. Dyslin. **Conflict of Interest Statement:** The authors declare no competing financial interests.

## Abstract

Oral reading relies on lexical and sublexical processes with distinct neural mechanisms. Damage within the sublexical system causes phonological alexia, a blanket diagnosis describing acquired deficits in reading unfamiliar words. Improving the precision of alexia diagnosis requires understanding the neurocognitive basis of specific reading subprocesses. This study investigated the neural correlates of sublexical reading in 64 adults with chronic left-hemisphere stroke (LHS), focusing on lesions that impair the use of learned orthography-to-phonology (OP) mappings to read new words. Participants read aloud real words and three types of pseudowords varying in the number of plausible OP mappings at the level of the orthographic body: zero mappings (0M), one mapping (1M), and multiple mappings (MM). LHS participants exhibited phonological reading deficits with an exaggerated lexicality effect compared to 71 neurotypical controls. Across both groups, pseudowords with learned OP mappings were read more accurately than those without. Voxelwise and connectome-based lesion-symptom mapping revealed that relative lexical reading deficits were associated with lateral temporal lesions, while sublexical reading deficits were associated with lesions or disconnections of the left inferior frontal (IFG), supramarginal, and pre/postcentral gyri. Applying learned OP mappings relied on anterior IFG and frontoparietal connections, while resolving multiple plausible OP mappings relied on intraparietal connections. These results underscore the role of learned mutigraphemic OP mappings in sublexical reading, and demonstrate that disruptions of different sublexical reading subprocesses result in subtly different deficit patterns. Dissecting the neurocognitive basis of reading subprocesses may improve the precision of alexia diagnosis and point to new treatments.

## Introduction

Reading is an acquired skill that is fundamental for engagement in contemporary society. Among individuals who experience language impairments following left-hemisphere stroke, approximately two-thirds report specific challenges with reading, known as alexia, which significantly diminish independence and quality of life (Brookshire et al., 2014; Purdy et al., 2019). To better understand alexia and contribute to improved diagnosis and treatment planning, it is essential to understand both the cognitive and neural bases of reading.

Reading hinges on the dynamic interplay among three key types of knowledge: orthography (i.e., spelling), phonology (i.e., sound), and semantics (i.e., meaning) of written words. Central to all leading cognitive models of reading is the recognition of at least two pathways facilitating successful oral reading (Coltheart, 2006; Coltheart et al., 2001; Perry et al., 2007; Plaut et al., 1996). The lexical pathway supports the accurate pronunciation of known words, relying on whole-word mappings of orthography to phonology, potentially influenced by associated semantic content. Conversely, the sublexical pathway provides a direct link from orthography to phonology, aiding the decoding of letter combinations to their corresponding sounds. This pathway facilitates the pronunciation of novel words and pseudowords which lack stored lexical and semantic content. Compelling evidence for the distinction between these pathways emerges from alexia research, where dissociable syndromic patterns of alexia correspond to the lexical and sublexical routes. For example, phonological alexia is defined by a selective impairment of pseudoword reading relative to word reading that has been interpreted as an impairment to the sublexical pathway in the framework of reading models (Beauvois & Derouesne, 1979).

Prior research into the mechanisms underlying sublexical reading has proposed that readers can either determine the pronunciation of novel words through orthography-to-phonology (OP) decoding or by using analogies to known words, which may involve indirect activation of the lexical-semantic knowledge (Andrews & Scarratt, 1998; Patterson & Lambon Ralph, 1999). Given English’s orthographic inconsistencies, the letter groups that correspond to the orthographic body or syllable rime (vowel nucleus plus an optional consonant cluster in the coda) are especially utilized in spelling-to-sound translation (Bowey, 1990; Treiman et al., 1995). Careful orthotactic manipulations at the level of these sub-word OP mappings, specifically of the orthographic body, can therefore be useful to isolate mechanisms involved during different types of sublexical reading. For example, the orthographic body of pseudowords can be regular with only one plausible OP mapping (1M pseudowords; e.g., ‘bink’ pronounced like ‘wink’), irregular with multiple plausible OP mappings (MM pseudowords; e.g., ‘tave’ which can be pronounced like ‘have’ or ‘gave’), or unique with no analogous orthographic bodies in English (0M pseudowords; e.g., ‘dofe’) (Andrews & Scarratt, 1998; Taft, 1982). It remains unclear to what extent these mappings at the level of the orthographic body (hereafter referred to as learned OP mappings) facilitate oral reading of novel words, and whether different types of OP mappings rely on distinct cognitive and neural substrates. Based on this framework, reading of 1M and MM pseudowords should rely on learned mappings at the level of the orthographic body, followed by assembly with the orthographic onset into a plausible pronunciation of the pseudoword. In contrast, 0M pseudowords without orthographic body neighbors should rely only on learned mappings of grapheme sequences at a smaller grain size, followed by assembly of those smaller units. Research on oral pseudoword reading has also suggested a consistency effect in which MM pseudowords may elicit slower reading speeds compared to 1M pseudowords (Glushko, 1979; Taraban & McClelland, 1987), potentially due to the supplemental involvement of semantic and lexical processes or additional cognitive demand required to disambiguate between multiple plausible pronunciations. This may reflect the influence of frequency-weighted probabilistic production mechanisms within the sublexical route (Zevin & Seidenberg, 2006), underscoring the need for further research to clarify these processes and their neural underpinnings.

Understanding the neural substrates that support these cognitive processes is crucial for advancing our knowledge of reading mechanisms. Functional neuroimaging studies, primarily in neurotypical adults, have delineated the general neural organization of the reading network, which includes a dorsal, phonological pathway facilitating assembly and articulation during reading, and a ventral, lexical-semantic pathway enabling access to lexical knowledge including word meanings (Coltheart et al., 2001; Dehaene, 2009; Taylor et al., 2013). These pathways generally correspond to the sublexical and lexical pathways described in cognitive models of reading (Taylor et al., 2013). Lesion studies following left-hemisphere stroke have further corroborated this distinction, highlighting correlations between lesion locations and deficits within each pathway (Coltheart et al., 2001; Dickens et al., 2019; Taylor et al., 2013). The dorsal sublexical pathway is typically isolated using a lexicality effect, comparing pseudoword reading to real word reading, either in terms of increased activity during functional magnetic resonance imaging (fMRI) studies or behavioral deficits in lesion studies. This contrast is effective in isolating sublexical processes, as it controls for speech production and subtracts out lexical-semantic processing, thus highlighting mechanisms unique to sublexical processing.

In comparison to real word reading, pseudoword reading elicits greater activation of several key brain regions, including the left posterior fusiform gyrus, left inferior parietal cortex, left inferior frontal gyrus (IFG), left precentral gyrus, and left insula (Binder et al., 2003; Carreiras et al., 2007; Dietz et al., 2005; Fiez et al., 1999; Graves et al., 2010; Hagoort et al., 1999; Mechelli et al., 2003, 2007; Taylor et al., 2013). Researchers frequently attribute OP translation to the inferior parietal cortex, as it consistently shows increased activity for pseudoword reading tasks, spelling and rhyming tasks, and training on novel symbol-phoneme associations (Baldo & Dronkers, 2006; Hashimoto & Sakai, 2004). The left IFG, insular cortex, and precentral gyrus have also been implicated in phonological processing above and beyond general executive demands that are not specific to reading (Binder et al., 2005; Mechelli et al., 2007; Poldrack et al., 1999). Specifically, the left IFG and insula have been proposed to correspond with related but different components of the cognitive models, namely the phoneme system of the DRC, the phonological output buffer within CDP+ models, and the phoneme units within the triangle model (Taylor et al., 2013). A recent study of individuals with post-stroke alexia isolated brain regions corresponding to two different types of phonological impairment across several tasks, providing evidence for subspecialized regions within the sublexical route, with the inferior parietal cortex involved in sensorimotor translation and the ventral premotor cortex in motor phonology (Dickens et al., 2021). Thus, functional imaging and lesion studies have helped to delineate the types of phonological processing performed by different processing regions within the dorsal sublexical processing stream. Despite behavioral evidence that learned OP mappings at the level of the orthographic body are important for sublexical reading, the neural substrates underlying the use of these mappings for sublexical reading have not been examined.

In the current study, we aimed to bridge the gap in understanding the neural mechanisms of sublexical reading by 1) replicating and extending previous findings that lesions and disconnections in the dorsal phonological pathway result in sublexical reading deficits, and 2) isolating the specific regions within the sublexical pathway that facilitate the application of learned OP mappings during pseudoword reading. To achieve this, left-hemisphere stroke (LHS) survivors and demographically matched control subjects completed oral reading tasks involving both real words and pseudowords manipulated according to the number of learned OP mappings at the level of the orthographic body. We compared behavioral performance across groups and stimulus types, and then used support vector regression voxel-based lesion-symptom mapping (SVR-VLSM) and connectome-based LSM (SVR-CLSM) analyses to identify lesion locations and disconnections associated with sublexical reading deficits. Specifically, we aimed to isolate lesions leading to deficits in the use of learned OP mappings during sublexical reading, selection among multiple OP mappings, and sublexical reading in the absence of learned OP mappings at the level of the orthographic body. Our findings provide critical evidence for distinct neural substrates underlying sublexical and lexical reading, and that learned OP mappings facilitate sublexical reading via dissociable neural subprocessors within the sublexical pathway.

## Materials and Methods

### Participants

Participants included 64 adults with a history of chronic left-hemisphere stroke (LHS) and 71 neurotypical controls (see Table 1). Participants were native English speakers with no significant neurological or neuropsychiatric disorders other than stroke. Control participants with no history of stroke were matched to the stroke group on age and education. All participants provided written informed consent, and the Georgetown University Institutional Review Board approved the study protocol. Participants were recruited as a part of a larger study on post-stroke language outcomes (Clinicaltrials.gov NCT04991519). LHS participants were excluded if they could not complete the oral reading tasks (at least 5/200 real words). Years of education were determined based on the highest degree-level attained in the United States (high school = 12, college = 16, master’s = 18, JD = 19, MD = 20, PhD = 21).

**Table 1.**
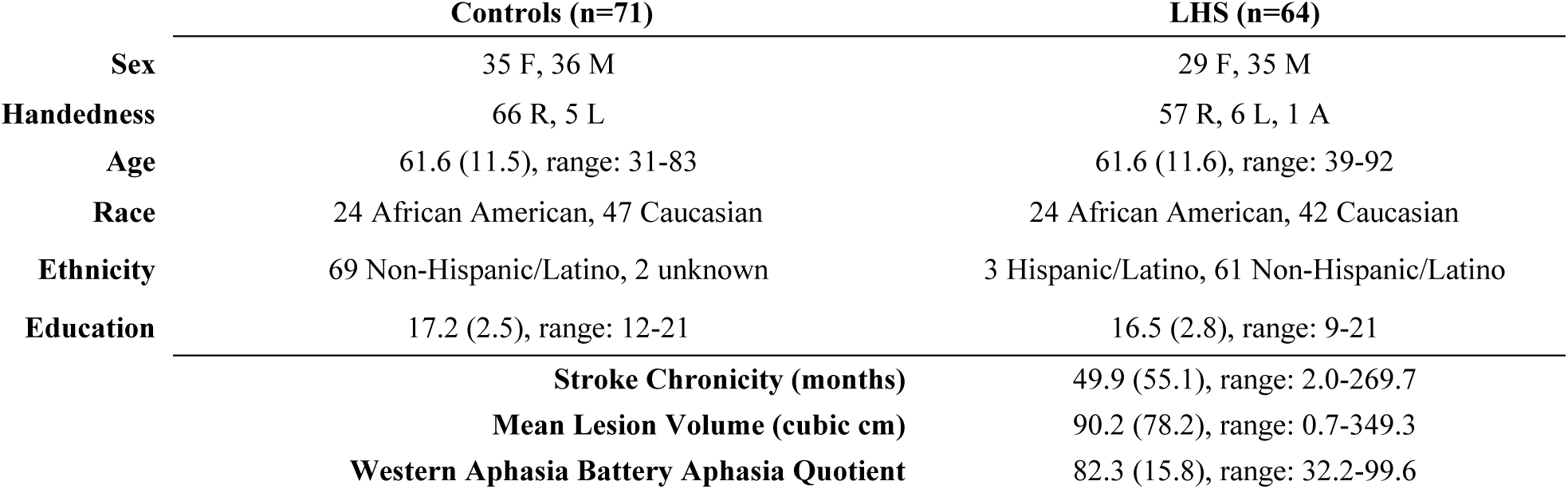
Participant Characteristics. Demographic information is listed for both groups as well as stroke and aphasia characteristics for left hemisphere stroke (LHS) participants. Handedness is self-reported as left (L), right (R), or ambidextrous (A).

### Oral Reading Assessments

Participants read aloud a list of 200 real words and 60 pseudowords. Real word oral reading included 200 monosyllabic words that varied orthogonally in frequency, spelling-to-sound regularity, and imageability. Three types of pseudowords were equally presented (20 stimuli each), differentiated based on the number of orthography-to-phonology body mappings that exist in English: zero mappings (0M), one mapping (1M) and multiple mappings (MM). 0M pseudowords (e.g., “dofe”) are pseudowords whose orthographic bodies do not exist within the English lexicon. 1M pseudowords (e.g., “bink”) have only one plausible pronunciation based on orthographic bodies in English (i.e., -ink as in “sink”). MM pseudowords (e.g., “chead”) involve at least two plausible pronunciations based on existing orthographic bodies in English (i.e., -ead pronounced either as in “bead” or “head”). Pseudoword stimuli was adapted and extended from Friedman et al. (1992, Supplemental Table 2). Pseudowords were matched to words on body consistency, positional bigram frequency, and articulatory complexity (Stoel-Gammon, 2010). All stimuli were monosyllabic and 3-6 letters in length.

Participants were seated at a table in a quiet testing room with a 17” Dell Inspiron touch-screen laptop in tent-mode. The tasks were administered in E-Prime 3.0. Real words were presented in blocks of 25 items with breaks between blocks. Pseudowords were presented in blocks of 20 items. The tasks were self-paced, and each trial timed out if the participant did not advance to the next item after 10 seconds. Responses were video recorded and scored offline. Scoring was completed based on the first complete reading attempt containing both a consonant and a vowel, and any response that included a plausible pronunciation of the presented pseudoword or real word was scored as correct. Responses to pseudowords were coded as incorrect if they contained spelling-to-sound mappings that do not occur in American English.

### Behavioral Analyses

Two logistic mixed effect models were estimated to determine the effects of group (LHS vs controls), lexicality, and pseudoword type on reading accuracy. The first model assessed the main effects of group and lexicality on reading accuracy, as well as their interaction (group by lexicality), while including fixed effects of age and education level and random effects of subject and item. The second model assessed the main effects of group and pseudoword type on pseudoword reading accuracy, as well as their interaction (group by pseudoword type), while including fixed effects of age and education level and random effects of subject and item. Pseudoword type was dummy coded within the model with 0M pseudowords serving as the reference category.

Four additional mixed effects models were estimated to determine the effects of lexicality and pseudoword type on reading accuracy within each group with the same fixed effects of age and education level and random effects of subject and item. All models were fitted using the *glmer* function in R Statistical Software with maximum likelihood estimation (v4.1.2; R Core Team 2021). Significance levels were determined at p < 0.05. Results of the logistic mixed effects model are reported as odds ratios and associated 95% confidence intervals, with statistical significance determined through the Wald Z-statistic and its associated p-value.

### Neuroimaging

#### Structural imaging

All brain images were acquired on Georgetown University’s 3T Siemens Prisma scanner. Scans of each participant included a T1-weighted magnetization prepared rapid gradient echo (MPRAGE), a fluid-attenuated inversion recovery (FLAIR) sequence, and diffusion imaging sequence. High resolution T1-weighted images were acquired for all participants with the following parameters: 176 sagittal slices; slice thickness = 1 mm, field of view = 212 x 212 mm; matrix = 256 x 256; repetition time (TR) = 300 ms; echo time (TE) = 6 ms; 1 mm cubic voxels. The FLAIR sequence aided in manual lesion tracing and was acquired with the following parameters: 192 sagittal slices, slice thickness = 1 mm; 1 mm^3^ voxels; flip angle = 120°; FOV = 256 mm, matrix = 256 mm x 256; TR = 5000 ms; TE = 386 ms; Inversion Time = 1800 ms; scan time: ∼ 5 mins. Diffusion Imaging scans were acquired to quantify white matter connections at these parameters: HARDI; TR = 5.0 s, TE = 0.082 s, readout time = 0.061 s, diffusion-weighted gradients: 81 directions at b = 3000, 40 at b = 1200, 7 at b = 0, 70 slices, 2 mm cubic voxels, flip angle = 90, phase encoding direction = anterior to posterior, partial Fourier = 6/8, field of view (FOV) = 232 mm, matrix = 116 x 116, slice acceleration = 1.

#### Lesion tracing and normalization

Stroke lesions were manually segmented on each participant’s MPRAGE and FLAIR images using ITK-SNAP software and approved by P.E.T. MPRAGEs and lesion masks were warped to the Clinical Toolbox Older Adult Template using Advanced Normalization Tools through a custom standard pipeline described elsewhere (Avants et al., 2011; K. C. Martin et al., 2024) (ANTs; http://stnava.github.io/ANTs/).

#### Structural connectome construction

Structural connectomes were derived from multi-shell high angular resolution diffusion imaging (MS-HARDI) scans according to methods described in Dickens et al. (2021). Structural connectivity was quantified through 15 million streamlines generated by probabilistic anatomically constrained tractography (Smith et al., 2012) on the white matter fiber orientation distributions in native space (algorithm = iFOD2, step = 1, min/max length = 10/300, angle = 45, backtracking allowed, dynamic seeding, streamlines cropped at grey matter-white matter interface). Edges of the structural connectome were generated by assigning streamlines to parcels of the Lausanne atlas at scale 125 (https://github.com/mattcieslak/easy_lausanne) (Daducci et al., 2012), yielding a 2D structural connectivity weighted adjacency matrix for each participant. Each edge in the connectome is proportional to the cross-sectional area of white matter connecting the two parcels at the intersecting row-column. We then create a disconnection matrix for each stroke survivor by labeling each connection between brain parcels (i.e., an edge) in the stroke survivors’ connectomes as lesioned if it was absent in a stroke survivor’s connectome but present in 100% of control subjects’ connectomes. This procedure yielded a binary disconnection matrix for each stroke survivor in which disconnected edges have a value of 1 and intact connections have a value of 0, analogous to a binary voxelwise lesion mask.

#### Lesion overlap maps

The overlap of the patients’ lesions reveals widespread damage predominantly in the perisylvian area, characteristic of strokes of the left middle cerebral artery (Fig. 1a). The disconnectome overlap map shows a significant loss of interhemispheric connections and connections within the left hemisphere as expected (Fig. 1b)

**Figure 1.**
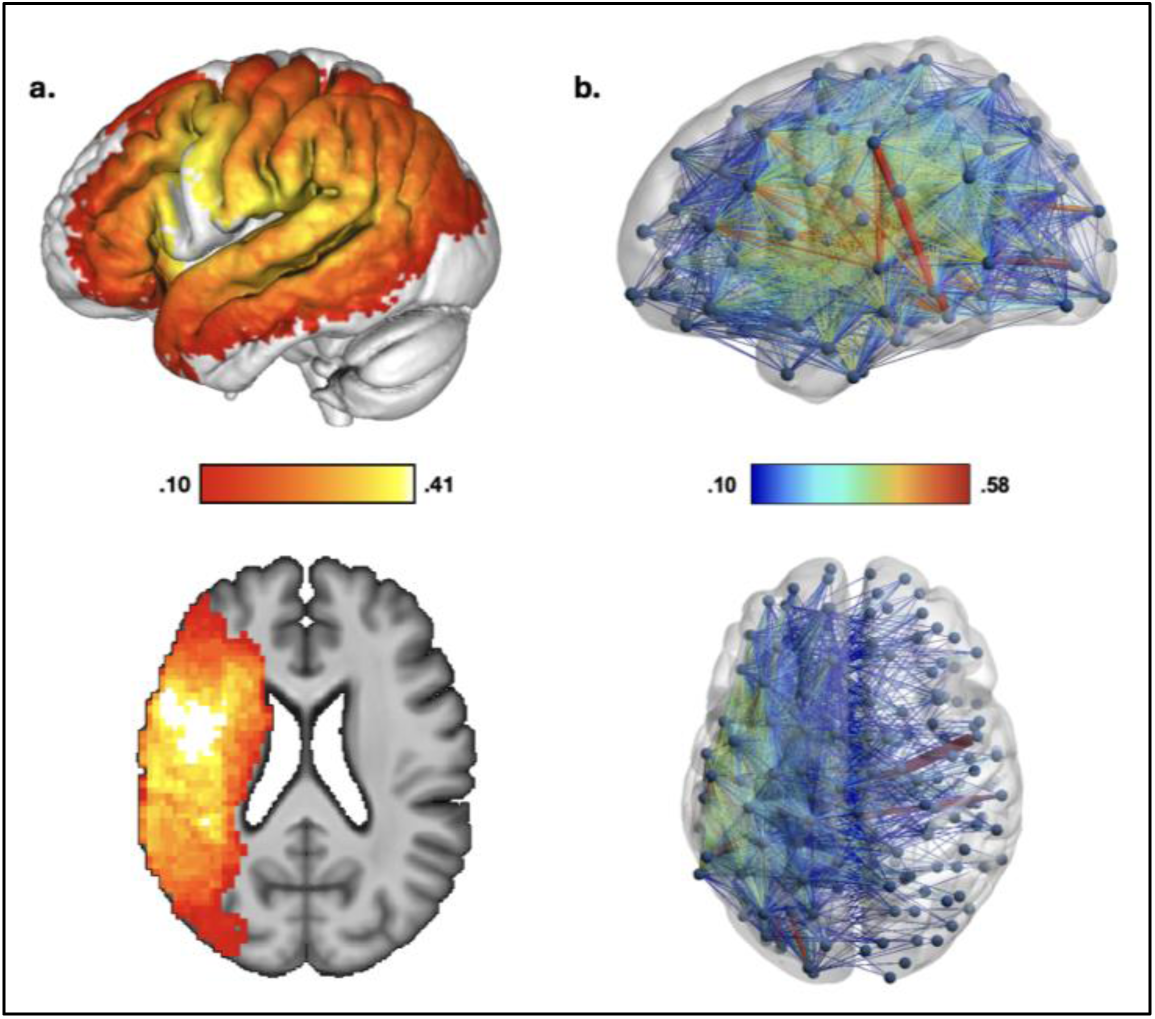
Voxel and Connectome-based lesion overlaps. Lesion overlap maps are shown on voxelwise **(a)** and connectome **(b)** levels for the 64 LHS participants. Color scales indicate the number and proportion of participants with lesions to those voxels and disconnections.

**Table 2.** Descriptive Statistics for Oral Reading Accuracy in Controls and LHS Participants.

|  | Real Words<br>(n=200) |  | All Pseudowords<br>(PW; n=60) |  | 0M PWs<br>(n=20) |  | 1M PWs<br>(n=20) |  | MM PWs<br>(n=20) |  |
| --- | --- | --- | --- | --- | --- | --- | --- | --- | --- | --- |
|  | Controls | LHS | Controls | LHS | Controls | LHS | Controls | LHS | Controls | LHS |
| Missing | 3 | 1 | 1 | 0 | 1 | 0 | 1 | 0 | 1 | 0 |
| Mean | 94.3% | 75.0% | 93.0% | 48.9% | 89.7% | 41.2% | 94.3% | 52.9% | 93.2% | 52.7% |
| Std. Dev. | 12% | 24% | 15% | 31% | 16% | 30% | 15% | 34% | 14% | 33% |
| Min. | 77.5% | 0.09% | 22% | 0% | 20% | 0% | 15% | 0% | 25% | 0% |
| Max. | 100% | 98.5% | 100% | 100% | 100% | 100% | 100% | 100% | 100% | 100% |

### Lesion-symptom mapping analyses

To identify the brain regions subserving reading and sublexical processes facilitating the application of learned OP mappings, we conducted support vector regression lesion-symptom mapping (SVR-LSM) at the voxelwise and connectome levels (Zhang et al., 2014). The SVR-VLSM analyses were conducted using a MATLAB toolbox developed by our group (DeMarco & Turkeltaub, 2018). Analyses were limited to voxels lesioned in at least 10% of participants. Covariates of lesion volume, age, and education were regressed out of both the lesion and behavioral data prior to modeling.

Our SVR-CLSM approach is an extension of SVR-VLSM which quantifies lesion through anatomical disconnection and thus identifies parcel-based neuroanatomical networks, as opposed to voxel-based clusters. The SVR-CLSM analyses complement the SVR-VLSM analyses by identifying the necessary contributions of both lesioned and spared, but disconnected, brain regions to reading. Lesion volume, age, and education were regressed out of both the connectome edge values and behavioral data. Only edges at which greater than 10% of stroke participants had lesions (i.e. 0 edges after binarization) were included in the analysis. Connections not present in 100% of control subjects were excluded from analyses in order to reduce Type I error. Statistical significance of the SVR-CLSM beta-maps was determined via a permutation-based FWER correction [10 000 permutations, FWER P < 0.05, one-tailed (negative)] (Kimberg et al., 2007) and was evaluated at both the edge-level (disconnection between pairs of brain parcels) and parcel-level (disconnection of a single brain parcel, including all anatomical endpoints).

We conducted 5 pairs of voxel-based and connectome-based SVR-LSM analyses to isolate lesion locations and disconnections associated with specific reading and sublexical processing deficits. First, we identified brain regions that are important for sublexical reading with SVR-LSM of accuracy on all pseudowords controlling for accuracy on all real words. This first contrast identifies lesion locations associated with an exaggerated lexicality effect (consistent with diagnoses of phonological alexia). Next, SVR-LSM was conducted for the reverse contrast, accuracy on all real words controlling for pseudowords, to identify lesions associated with the reduction of a lexicality effect.

Our next three critical analyses elaborated on sublexical processes. The first sublexical contrast aimed to isolate neural correlates important for the application of learned OP mapping during sublexical reading, by identifying lesion locations and disconnections associated with reading accuracy on pseudowords with existing OP mappings (MM and 1M) while covarying for two variables: accuracy on pseudowords with zero mappings (0M) and accuracy on real words. The second sublexical contrast aimed to isolate neural correlates underlying the sublexical processes important for accuracy on pseudowords with multiple plausible OP mappings (MM) covarying for 1M pseudowords and accuracy on real words. The final sublexical contrast aimed to isolate neural correlates important for sublexical processes given unfamiliar orthographic bodies and therefore absent of relevant learned OP mappings, specifically accuracy on 0M pseudowords controlling for accuracy on MM and 1M pseudowords and accuracy on real words.

## Results

### Behavioral Results

LHS participants showed greater variability in performance than controls participants, as expected (Table 2, Figure 2). The first logistic mixed effect model showed an expected main effect of group (Z = -8.91, P = < 0.001, OR = 0.070, 95% CI = 0.04-0.13) such that control participants read more accurately overall compared to LHS participants (Table 3). There was also a main effect of lexicality (Z = -5.65, P = < 0.001, OR = 0.337, 95% CI = 0.23-0.49) indicating that accuracy across groups was significantly higher for words compared to pseudowords. A significant interaction of group and lexicality was observed (Z = -8.79, P = < 0.001, OR = 0.398, 95% CI = 0.32-0.49), demonstrating an exaggerated lexicality effect, i.e., phonological reading deficits, in left-hemisphere stroke participants overall relative to controls.

**Figure 2.**
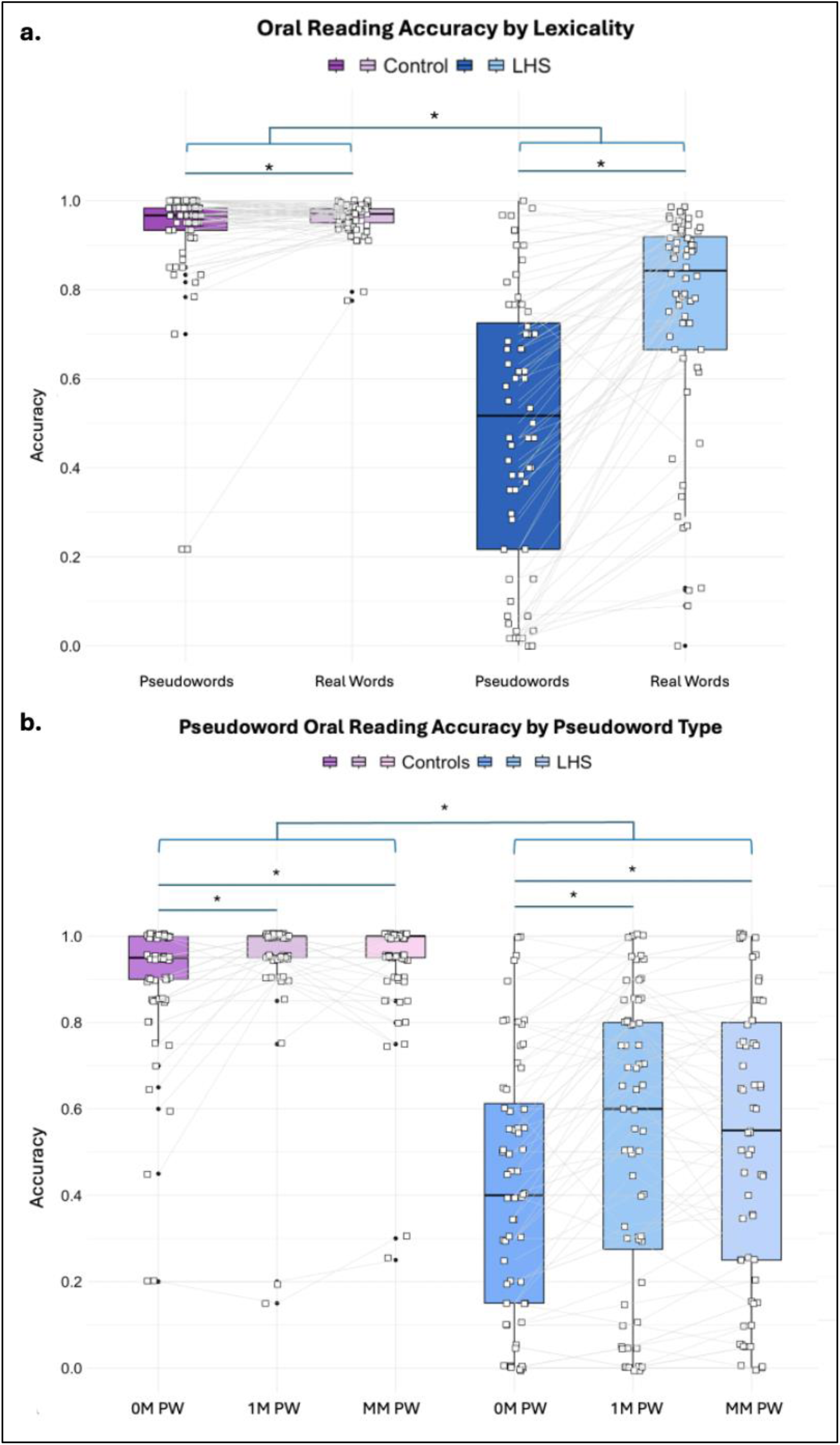
Oral reading accuracy by lexicality and pseudoword type. **(a)** The average accuracies on responses across trials for real word oral reading and pseudoword oral reading which contains all pseudoword types are shown for control participants (left) and LHS participants (right). White squares denote individual participants, and gray lines connect the same individual’s average performance on the two tasks. Each embedded boxplot has a bold black line designating the mean accuracy for each task and participant group. Solid lines above the data distributions indicate significant main effects of group, lexicality (2a), and PW type (2b). See Table 2 for descriptive statistics and Tables 3-6 for results of the mixed effect models. **(b)** The average accuracies on responses across trials for each pseudoword type within the oral pseudoword reading task are shown for control participants (left) and LHS participants (right).

**Table 3.** Results of a model examining lexicality and history of stroke to accuracy reading aloud.

Formula: Accuracy ~ 1+ Education by degree + Age + Group\*Lexicality + (1|Subject) + (1|Item)
| Predictors | Odds Ratios | CI | p | p<0.05 |
| --- | --- | --- | --- | --- |
| Education | 1.082 | 0.97-1.21 | 0.159 |  |
| Age | 1.015 | 0.99-1.04 | 0.255 |  |
| Group | 0.070 | 0.04-0.13 | <0.001 | * |
| Lexicality | 0.337 | 0.23-0.49 | <0.001 | * |
| Group*Lexicality | 0.398 | 0.32-0.49 | <0.001 | * |
| **Random Effects** |  |  |  |  |
| Random Intercept: Item | 1.30 |  |  |  |
| Random Intercept: Subject | 2.76 |  |  |  |
| N<sub>subject</sub> | 135 |  |  |  |
| N<sub>item</sub> | 260 |  |  |  |

If learned OP mappings facilitate sublexical reading, then both groups should show increased accuracy for pseudowords with learned mappings (1M and MM pseudowords) relative to those without any orthographic body neighbors in English (0M pseudowords). Consistent with this hypothesis, a logistic mixed effect model revealed a significant main effect of pseudoword type on pseudoword reading accuracy, such that across groups 1M and MM pseudowords were read with increased accuracy relative to 0M pseudowords (Z = 3.63, P = < 0.001, OR = 2.58, 95% CI = 1.55-4.29; Z = 2.54, P = 0.011, OR = 1.90, 95% CI = 1.16-3.12 respectively). This model also showed a significant main effect of group such that left hemisphere stroke was related to reduced pseudoword reading accuracy (Z = -10.48, P = < 0.001, OR = 0.337, 95% CI = 0.23-0.49). There were no significant interactions of group by pseudoword type. Full model estimates are shown in Table 4.

**Table 4.** Results of a model relating pseudoword type and history of stroke to accuracy reading aloud.

Formula: Accuracy ~ 1+ Education by degree + Age + Group\*Pseudoword Type + (1|Subject) + (1|Item)
| Predictors | Odds Ratios | CI | p | p<0.05 |
| --- | --- | --- | --- | --- |
| Education | 1.12 | 0.98-1.28 | 0.094 |  |
| Age | 1.00 | 0.97-1.03 | 0.784 |  |
| Group | 0.021 | 0.01-0.04 | <0.001 | * |
| 1M Pseudowords | 2.58 | 1.55-4.29 | <0.001 | * |
| MM Pseudowords | 1.90 | 1.16-3.12 | <0.011 | * |
| Group * 1M Pseudowords | 0.85 | 0.57-1.27 | 0.422 |  |
| Group * MM Pseudowords | 1.13 | 0.77-1.66 | 0.541 |  |
| **Random Effects** |  |  |  |  |
| Random Intercept: Item | 0.36 |  |  |  |
| Random Intercept: Subject | 3.64 |  |  |  |
| N<sub>subject</sub> | 133 |  |  |  |
| N<sub>item</sub> | 60 |  |  |  |

Additional within-group mixed effect models were estimated to determine effects of lexicality and pseudoword type without group as a fixed effect (see Table 5 and 6). A main effect of lexicality was observed in both control and LHS participants such that real words were read with greater accuracy than pseudowords (controls: Z = -5.84, P = < 0.001, OR = 0.165, 95% CI = 0.09-0.30; LHS participants: Z = -12.19, P = < 0.001, OR = 0.143, 95% CI = 0.10-0.20). A main effect of pseudoword type was also observed in both control and LHS participants for both 1M and MM pseudowords relative to 0M pseudowords (see Table 6). Both models in controls also revealed a significant main effect of education such that more prior education related to increased reading accuracy (see Table 5 and 6). The lexicality effect model in LHS participants revealed a significant main effect of age such that increased age related to higher reading accuracy (see Table 5 and 6).

**Table 5.** Results of two models relating lexicality to accuracy reading aloud within control participants (5a) and LHS participants (5b).

a. Control Participants -- Formula: Accuracy ~ 1+ Education by degree + Age + Lexicality + (1|Subject) + (1|Item)
| Predictors | Odds Ratios | CI | p | p<0.05 |
| --- | --- | --- | --- | --- |
| Education | 1.218 | 1.08-1.38 | 0.002 | * |
| Age | 0.989 | 0.96-1.02 | 0.384 |  |
| Lexicality | 0.165 | 0.09-0.30 | <0.001 | * |

| Random Effects |  |
| --- | --- |
| Random Intercept: Item | 3.25 |
| Random Intercept: Subject | 1.41 |
| N<sub>subject</sub> | 70 |
| N<sub>item</sub> | 260 |

b. LHS participants -- Formula: Accuracy ~ 1+ Education by degree + Age + Lexicality + (1|Subject) + (1|Item)
| Predictors | Odds Ratios | CI | p | p<0.05 |
| --- | --- | --- | --- | --- |
| Education | 0.947 | 0.79-1.13 | 0.548 |  |
| Age | 1.048 | 1.00-1.09 | 0.029 | * |
| Lexicality | 0.143 | 0.10-0.20 | <0.001 | * |
| Random Effects |  |  |  |  |
| Random Intercept: Item | 1.04 |  |  |  |
| Random Intercept: Subject | 3.73 |  |  |  |
| N<sub>subject</sub> | 65 |  |  |  |
| N<sub>item</sub> | 260 |  |  |  |

**Table 6.** Results of two models relating pseudoword type to accuracy reading aloud within control participants (6a) and LHS participants (6b).

a. Control participants -- Formula: Accuracy ~ 1+ Education by degree + Age + Pseudoword Type + (1|Subject) + (1|Item)
| Predictors | Odds Ratios | CI | p | p<0.05 |
| --- | --- | --- | --- | --- |
| Education | 1.320 | 1.14-1.53 | <0.001 | * |
| Age | 0.982 | 0.98-1.01 | 0.265 |  |
| 1M Pseudowords | 2.651 | 1.54-4.57 | <0.001 | * |
| MM Pseudowords | 1.958 | 1.15-3.33 | <0.001 | * |
| Random Effects |  |  |  |  |
| Random Intercept: Item | 0.41 |  |  |  |
| Random Intercept: Subject | 1.74 |  |  |  |
| N<sub>subject</sub> | 69 |  |  |  |
| N<sub>item</sub> | 60 |  |  |  |

b. LHS participants -- Formula: Accuracy ~ 1+ Education by degree + Age + Pseudoword Type + (1|Subject) + (1|Item)
| Predictors | Odds Ratios | CI | p | p<0.05 |
| --- | --- | --- | --- | --- |
| Education | 0.945 | 0.77-1.16 | 0.593 |  |
| Age | 1.022 | 0.97-1.07 | 0.378 |  |
| 1M Pseudowords | 2.190 | 1.45-3.31 | <0.001 | * |
| MM Pseudowords | 2.155 | 1.43-3.26 | <0.001 | * |
| Random Effects |  |  |  |  |
| Random Intercept: Item | 0.34 |  |  |  |
| Random Intercept: Subject | 4.88 |  |  |  |
| N<sub>subject</sub> | 64 |  |  |  |
| N<sub>item</sub> | 60 |  |  |  |

### Lesion Symptom Mapping

*Lexicality contrasts.* Two SVR-LSM analyses were performed to investigate whether damage to specific brain regions predicted reduced oral reading accuracy of pseudowords and real words, each controlling for the other stimulus type. Lesions involving left supramarginal gyrus (SMG) related to reduced oral pseudoword reading accuracy (SVR-VLSM clusterwise P = 0.03; cluster size: 5.500 cm^3^; center of mass: -48.4, -30, 26; Fig. 3a). 53 disconnections similarly related to reduced oral pseudoword reading accuracy (SVR-CLSM edgewise FWER P < 0.05; Fig 3b, Supplemental Table 1). Of these 53 disconnections, 17 connected LH frontal and parietal regions, 17 were intraparietal connections, and four were interhemispheric connections. Several regions were particularly implicated by these findings, including the left postcentral gyrus, the left supramarginal gyrus, and the left middle frontal gyrus (involved in 31, 22, and 11 disconnections respectively). The left precentral gyrus, left inferior parietal lobule, and left superior temporal sulcus were also each involved in 6 disconnections. Lesions to the middle temporal gyrus, superior temporal sulcus, and surrounding cortex and white matter resulted in the relative reduction of oral real word reading accuracy (SVR-VLSM clusterwise P = 0.02; cluster size: 7.281 cm^3^; center of mass MNI coordinates: -56, -42.6, 6.5; Fig. 4). No significant disconnections were identified related to the relative reduction of oral real word reading accuracy.

**Figure 3.**
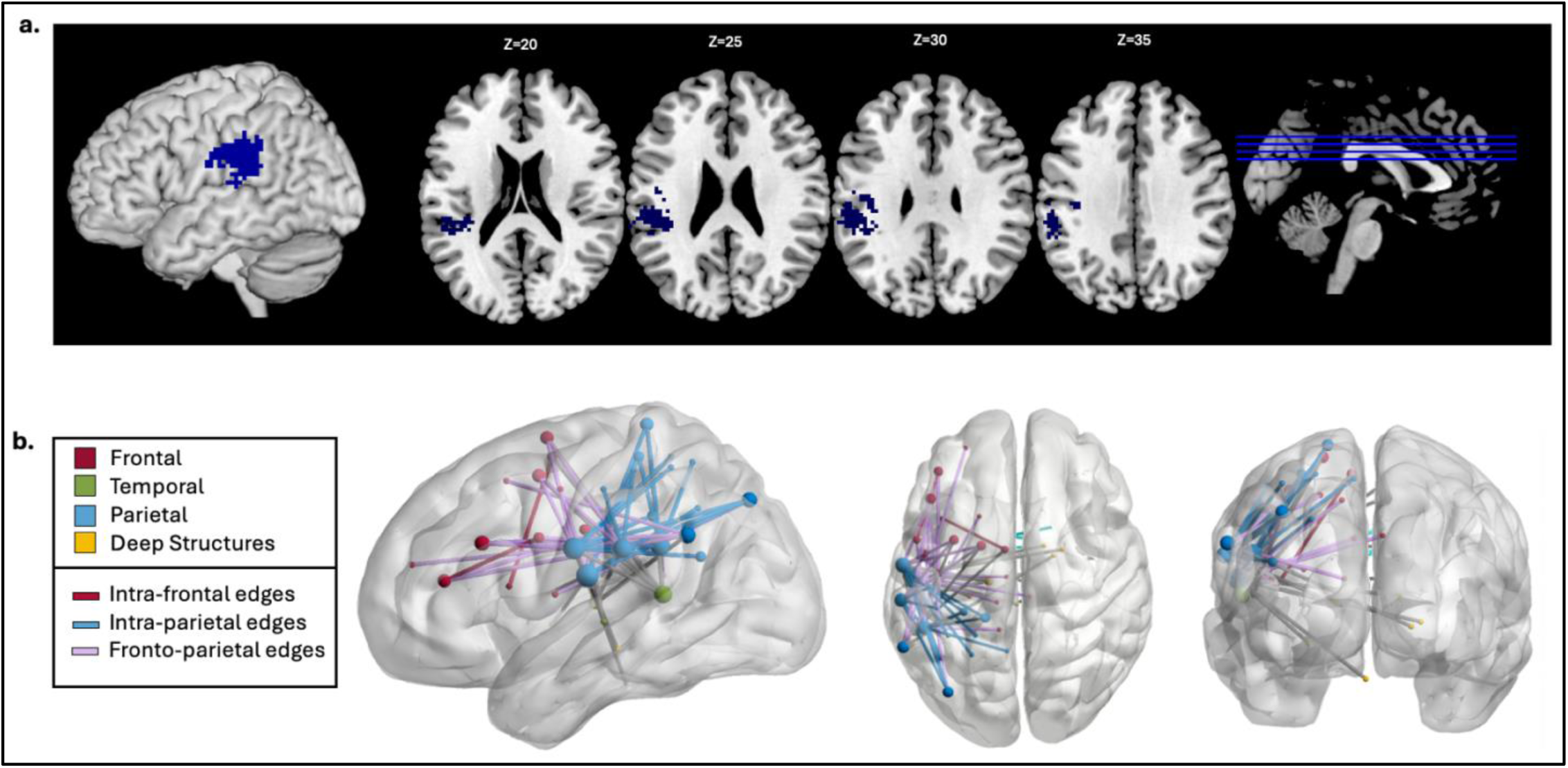
Lesion Symptom Mapping for Pseudowords > Real words. **(a)** Voxelwise LSM results at p<0.005 and clusterwise FWE p<0.05. **(b)** Connectome LSM results show significant edges color-coded by lobe. Node size relates to the number of significant edges at that node.

**Figure 4.**
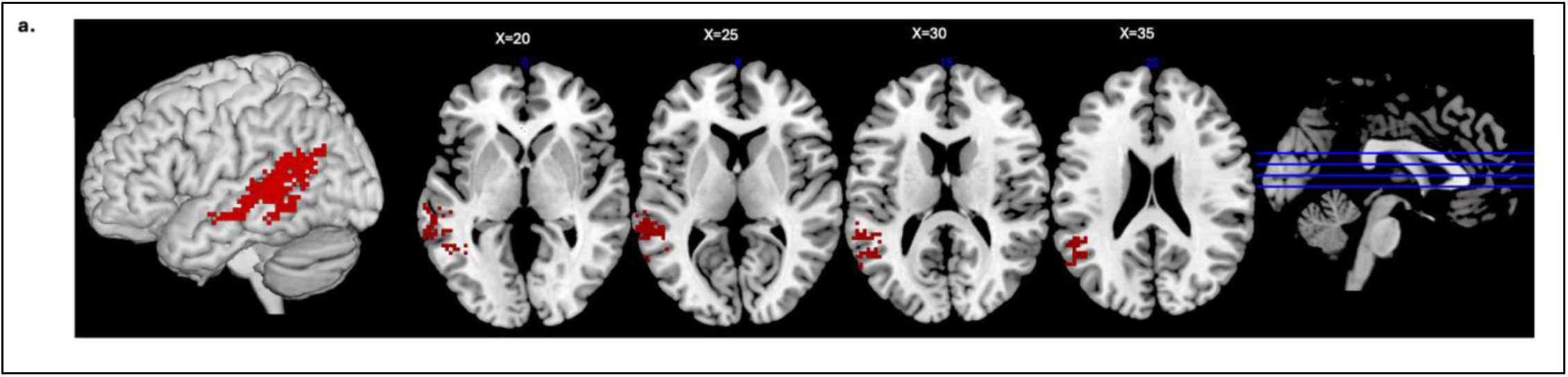
Lesion Symptom Mapping for Real Words > Pseudowords. Voxel-wise LSM results at p<0.005 and clusterwise FWE p<0.05.

#### Pseudoword contrasts by OP mapping

The first contrast of pseudoword types aimed to identify lesions associated with loss of the ability to use learned OP mappings for sublexical reading. Lesions to a cluster extending from the anterior inferior frontal gyrus, through the external capsule, and into to the ventral precentral gyrus resulted in relative reduction of accuracy on pseudowords with body-level OP mappings (MM + 1M), controlling for those without learned mappings in English (0M) (SVR-VLSM clusterwise P=0.01; cluster size: 8.109 cm3; center-of mass MNI coordinates: -37.7, 9.6, 12.4, Figure 5a). CLSM revealed 36 significant disconnections associated with reduced MM and 1M pseudoword reading (p*<*0.05, Figure 5b, Supplemental Table 1). Of these 36 disconnections, 14 connected LH frontal and parietal regions, 9 were LH intrafrontal connections, 5 were LH fronto-temporal connections, and 7 were interhemispheric connections. The regions associated with the highest number of significant disconnections included the middle frontal gyrus (19 connections), the precentral gyrus (15), the postcentral gyrus (11), and the SMG (7).

**Figure 5.**
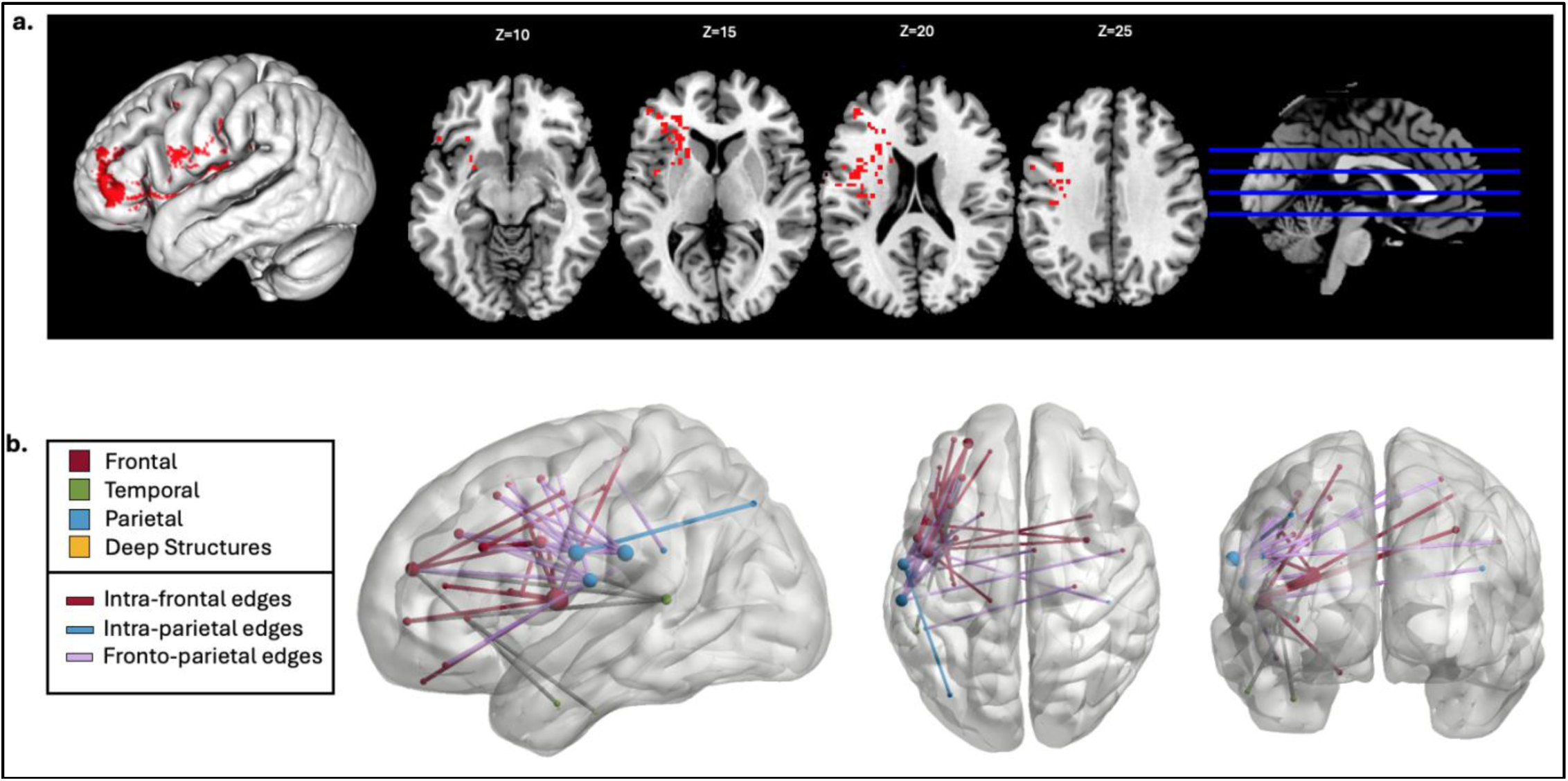
Lesion Symptom Mapping for 1M + MM Pseudowords > 0M Pseudowords. **(a)** Voxel-wise LSM results at p<0.005 and clusterwise FWE p<0.05. **(b)** Connectome LSM results show significant edges color-coded by lobe. Node size relates to the number of significant edges at that node.

Next, we aimed to identify lesions that result in impaired reading of pseudowords with multiple plausible OP mappings compared to those with single consistent mappings in English. Lesions to the anterior regions of the supramarginal gyrus were related to a relative reduction of MM pseudowords accuracy controlling for 1M accuracy but this result did not reach statistical significance (SVR-VLSM clusterwise p=0.058, cluster size: 3.625 cm3; center-of mass MNI coordinates: -35.6, -26.7, 20.2, Figure 6a). CLSM revealed 55 significant disconnections associated with deficits in MM pseudowords (p<.05, Figure 6b, Supplemental Table 1). These disconnections were primarily intraparietal and posterior frontoparietal in nature, including 14 intrafrontal, 13 frontoparietal, 8 parietal-subcortical, 3 fronto-subcortical, 2 temporoparietal, 1 frontotemporal, and 14 interhemispheric connections. Regions most frequently implicated included the postcentral gyrus (36 disconnections), precentral gyrus (9 disconnections), and supramarginal gyrus (13 disconnections).

**Figure 6.**
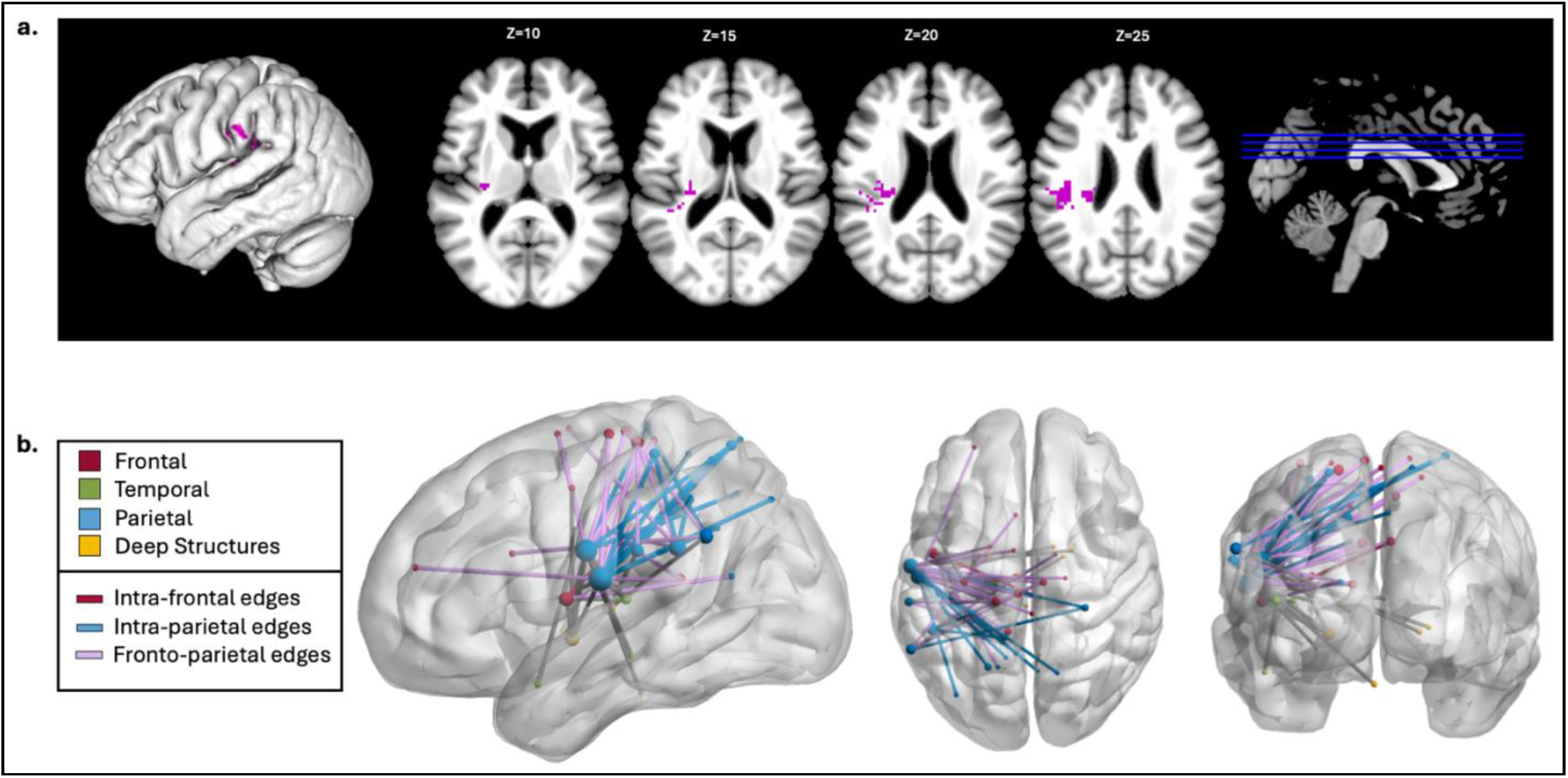
Lesion Symptom Mapping for MM Pseudowords > 1M Pseudowords. **(a)** Trending voxel-wise LSM results at p=0.058 and clusterwise FWE p<0.05. **(b)** Connectome LSM results show significant edges color-coded by lobe. Node size relates to the number of significant edges at that node.

Finally, we aimed to identify lesions associated with specific deficits in sublexical reading without learned OP mappings of orthographic bodies. No significant lesions or disconnections were identified for the third sublexical contrast, the relative reduction of reading pseudowords without learned mappings (0M), controlling for pseudowords with learned mappings (MM, 1M).

## Discussion

The primary aim of this study was to dissect the role of learned OP mappings in the sublexical route for reading. Our results clarify the distinction between the neuroanatomical bases of lexical and sublexical reading and affirms the role of multigraphemic learned OP mappings at the level of the orthographic body in sublexical reading. Depending on the nature of the orthographic body, reading an unfamiliar word relies on different subnetworks of processors within the broader frontoparietal sublexical reading network. These findings motivate further research into the mechanisms that underlie these subprocesses and the clinical relevance of deficits in applying learned OP mappings in alexia.

### Lesion evidence for sublexical and lexical routes for reading in the brain

Cognitive models of reading posit distinct processing routes for lexical and sublexical reading, particularly emphasizing that pseudowords, due to their lack of lexical content, are preferentially processed through the sublexical pathway (Coltheart et al., 2001; Perry et al., 2010; Plaut et al., 1996). Building upon prior activation and lesion studies suggesting that these reading pathways map onto distinct, distributed neural networks (Dickens et al., 2019; A. Martin et al., 2015; Price, 2012), our application of SVR-LSM to a large cohort of LHS survivors with variable reading impairments provides additional evidence that sublexical decoding of orthography-to-phonology relies predominantly on dorsal perisylvian regions, while lexical processing is more dependent on ventral areas surrounding the superior temporal gyrus.

The syndrome of phonological alexia, which is defined based on the presence of pseudoword reading deficits, has been associated with large and variable lesions throughout perisylvian regions (Rapcsak et al., 2009; Ripamonti et al., 2014). In two prior LSM studies, our group examined phonological reading deficits not based on syndromic classification, but rather based on the lexicality effect as a continuous measure, as we do here (Dickens et al., 2019, 2021). These studies found that lesions of ventral precentral gyrus and SMG resulted in phonological reading deficits, albeit with different characteristics. Ventral precentral gyrus lesions were associated with isolated difficulties with pseudoword reading, while SMG lesions were associated with preferential deficits on pseudowords along with milder deficits on matched real words. The participants for one of these studies partly overlapped with the participants examined here (Dickens et al., 2021), but both of the previous studies used a different stimulus set: 20 short pseudowords (3-5 letters), primarily with 1M bodies, and a set of real words differing by one letter. The current findings using a more complex set of pseudoword stimuli provide a replication of the main findings from these prior studies implicating the ventral precentral gyrus and SMG in sublexical reading.

The frontoparietal and intraparietal disconnections observed in our CLSM analysis implicate the wider dorsal processing network highlighted in prior research on sublexical reading (Aboud et al., 2016; Jobard et al., 2003; Taylor et al., 2013). Given that this analysis controls for accuracy in real word reading, the involvement of these regions is not solely related to processes such as letter recognition or speech production that are essential for all reading irrespective of lexical content. Consistent with the primary systems hypothesis (Woollams et al., 2018), our results identify disconnections involving regions subserving speech processing, including articulatory (left precentral gyrus), auditory (planum temporale), and somatosensory (ventral postcentral and supramarginal gyri) processing structures. Significant connections between the left postcentral gyrus, left SMG, left precentral gyrus, and left STS further align with the anatomy described in speech processing models (Bohland et al., 2010; Tourville & Guenther, 2011). The CLSM results implicated several nodes along the left middle frontal gyrus (MFG) and their connections to parietal regions, largely the SMG. These connections with the MFG may be responsible for aspects of executive function such as working memory and attentional control which contribute to performance on complex phonological tasks like oral pseudoword reading (Deschamps et al., 2014; Kim, 2020).

Regarding the lexico-semantic route, we found that lesions to the lateral temporal regions, centered on the superior temporal sulcus, cause a diminished lexicality advantage during oral reading (i.e., a reduction in the typical advantage for reading words compared to pseudowords). This finding corroborates previous functional imaging research localizing lexico-semantic processing to a ventral word reading pathway including the middle temporal gyrus (MTG) (Moore-Parks et al., 2010; Whitney et al., 2011). It also extends and aligns with prior lesion evidence identifying lesions to the posterior half of the middle temporal gyrus, extending into adjacent middle occipital gyrus resulting in regularization errors (Binder et al., 2016).

The lexicality advantage for reading can relate either to the use of whole word orthographic and phonological representations or to the use of semantic content to support word reading (Patterson & Lambon Ralph, 1999). To clarify the mechanisms underlying lexico-semantic reading, and perhaps to isolate subregions involved in application of different types of lexical knowledge for reading, future research should employ contrasts that assess factors such as imageability, frequency, part of speech, or morphosyntactic features. A relative sparing of real word reading compared to pseudoword reading after lesions to the SMG and fronto-parietal disconnections as found in our initial results suggests that the lexical pathway for reading does not pass through parietal regions. Instead, as suggested in some models of speech processing, the lexical pathway for reading may proceed anteriorly through the temporal lobe into inferior frontal cortex (Hickok, 2012). In this case, oral word reading may rely on direct connections between lexical representations in temporal cortex and articulatory plans for whole words in inferior frontal cortex.

### The role of OP mappings in sublexical reading

Our findings provide key insights into the mechanisms underlying sublexical reading by affirming the crucial role of learned OP mappings at the level of the orthographic body. Behaviorally, pseudowords containing orthographic bodies that are present in English words were read more accurately than those without analogous orthographic bodies in both LHS and control participants, demonstrating that readers can effectively apply familiar multigraphemic OP mappings to novel stimuli. Our behavioral data did not reveal a significant effect related to the presence of multiple learned OP mappings as compared to single mappings, suggesting that difficulty in pseudoword reading accuracy is more strongly influenced by the availability of learned OP mappings than by response ambiguity between competing mappings. Since we accepted any plausible pronunciation of the pseudowords as correct, this may have limited our ability to identify differences in performance related to the number of possible OP mappings. Effects of competing mappings might be observable in response times as originally noted in research assessing the effect of consistency on the reading speed of high and low frequency real words, and pseudowords (Glushko, 1979; Taraban & McClelland, 1987), though measures of accuracy remain more relevant for the evaluation of reading deficits after stroke since response time measurement is not typically feasible in clinical settings.

Our findings revealed involvement of the ventral precentral and post central gyri, SMG, anterior left IFG, and a network of fronto-parietal regions and connections in reading pseudowords with learned body-level OP mappings as compared to pseudowords without such mappings. The findings of all the analyses collectively implicate the postcentral and precentral gyri as highly interconnected regions supporting pseudoword reading relative to real word reading, particularly for pseudowords with learned body-level OP mappings. While these regions are well-known for their involvement in motor articulatory and somatosensory representations from models of speech production (Bohland et al., 2010; Hickok, 2012; Tourville & Guenther, 2011; Walker & Hickok, 2016), our results suggest a more nuanced role in reading. In a prior LSM study, we found that the pseudoword reading deficit caused by lesions to these structures was specifically associated with motor phonological impairments, which caused an isolated pseudoword reading deficit with no effect on matched real words (Dickens et al., 2021). Here, we found that ventral sensorimotor regions were particularly important for reading pseudowords with learned body-level OP mappings. The contribution of these structures to pseudoword reading cannot be explained purely by speech production deficits, since oral word reading performance was covaried in both the prior and the current study. Instead, the involvement of the precentral and postcentral gyri suggests that learning to read may co-opt speech motor and somatosensory units for functions beyond simple articulation (Dickens et al., 2021). Specifically, since the reading of pseudowords with learned OP mappings was disrupted after lesions to, and disconnections from, these regions, the precentral and postcentral gyri are likely involved in storing, accessing, or assembling learned OP mappings. This corresponds with the primary roles of these regions in storing and assembling motor plans for speech production. Like OP mappings, speech motor plans must link to sensory representations and are stored at various grain sizes depending on the frequency of use (Tourville & Guenther, 2011). Similar computations may underlie selection and assembly of speech motor plans and OP mappings at various grain sizes. As such, learning to read may coopt sensorimotor regions evolved for speech production to store and assemble learned OP mappings for reading unfamiliar words.

Our main lexicality results implicated the SMG in pseudoword reading overall, suggesting that the SMG plays a central role in sublexical reading with and without learned body-level OP mappings. This aligns with prior fMRI and TMS studies that have identified the SMG as a critical region involved with several types of phonological processing (Jobard et al., 2003; Oberhuber et al., 2016; Vigneau et al., 2006), including phonological working memory (Deschamps et al., 2014) and phonological decoding during reading (Jobard et al., 2003). A prior LSM study on lexicality effects in a partly overlapping cohort using different reading items implicated the SMG in sensorimotor translation processes important for both speech and reading (Dickens et al., 2021). In this prior study, sensorimotor translation deficits, as reflected by poor pseudoword repetition, related to alexia that preferentially affected pseudowords, but also real words with regular spellings, albeit to a lesser degree. Notably, in the current study, the MM>1M SVR-VLSM analysis also identified the SMG, suggesting it may be especially important for sublexical reading when multiple body-level OP mappings are available. The corresponding CLSM analysis implicated the left postcentral gyrus, left precentral gyrus, left SMG, and a broader network of intraparietal connections. This finding may indicate an increased reliance on learned OP mappings in the precentral and postcentral gyri, and phonological processing and working memory in the SMG and other posterior parietal regions when deciding between multiple plausible pronunciations for an unfamiliar word. Overall, it is clear that the SMG plays a central role in sublexical reading, but its exact role remains enigmatic. It is possible that subregions of the SMG perform different specific processes important to different aspects of phonology and sublexical reading, or that the SMG performs a computation that underlies various specific phonological operations and renders it important for sublexical reading in multiple ways. Disentangling the specific contributions of the SMG to sublexical reading and phonology should be a focus of future research.

The anterior IFG, which was implicated in reading pseudowords with learned OP mappings here (1M+MM pseudowords), has been previously implicated in selection and control processes, often linked to semantic retrieval (Binder et al., 2005; Mechelli et al., 2007; Poldrack et al., 1999; Taylor et al., 2013). The left IFG has been directly related to phonological processing, though typically in more dorsal and posterior regions such as the pars opercularis (Vigneau et al., 2006). In this context, one possible interpretation of the anterior IFG result here is that the IFG is engaged in suppressing potential lexicalizations or in drawing analogies between pseudowords and known real words or mappings. Alternatively, the IFG could provide a facilitatory control signal to boost OP mapping representations in sensorimotor regions or the SMG. This suggests that the anterior IFG may contribute to selecting appropriate phonological representations through either of these mechanisms. However, this interpretation is complicated by our second sublexical contrast, which did not implicate the anterior IFG in pseudowords with multiple OP mappings compared to those with a single learned OP mapping. If the IFG’s primary function were indeed in selecting relevant sublexical representations, we would expect greater reliance specifically in the sublexical contrast isolating pseudowords with multiple plausible pronunciations to select between (i.e., MM pseudowords). Therefore, our results suggest the IFG’s role is not in selecting representations during sublexical reading, but either in suppressing potential lexicalizations or accessing relevant lexical knowledge. These findings call for further investigation into the anterior IFG’s mechanistic role in sublexical processing.

The CLSM results also implicated connections to the left MFG in reading pseudowords with OP mappings vs. those without. The MFG has been implicated in representations of both orthographic and phonological information (Graves et al., 2023; Zhao et al., 2016) as well as in attentional and working memory demands related to language and phonological processing (Barbeau et al., 2023; Curtis, 2006; Mottaghy et al., 2003). Alternatively, studies of the gray matter correlates of working memory have suggested that the working memory loop for phonological content is more reliant on the SMG and posterior IFG while the anterior IFG, MFG, and angular gyrus (AG) are more related to semantic working memory (Horne et al., 2022; R. C. Martin & He, 2004; Paulesu et al., 1993; Vigneau et al., 2006). This framework allows for the possibility that mechanisms usually employed for semantic working memory processes in frontal regions may be recruited specifically when reading pseudowords with learned OP mappings, perhaps due to their associations with lexical items. Alternatively, the MFG may have more multi-modal, general processing capacities (Diachek et al., 2020). A number of interhemispheric connections were also implicated in reading using learned OP mappings. These connections involve subcortical and medial regions of the right hemisphere rather than regions homotopic to canonical language processing regions in the left hemisphere, again suggesting a general cognitive contribution to applying learned OP mappings while reading unfamiliar words.

Clinically, these results highlight the importance of developing a more nuanced diagnostic scheme for alexia. While a specific deficit of pseudoword reading relative to real word reading is the hallmark symptom of phonological alexia, this diagnosis encompasses a variety of specific processing deficits resulting in the common phenotype of impaired pseudoword reading. Like our prior study describing two types of phonological reading deficits related to different phonological subprocesses (Dickens et al., 2021), this study provides additional evidence that alexia may be better described based on specific process-level deficits. Further, these results highlight the potential of targeted rehabilitation approaches that leverage preserved neural circuits for phonological and lexical processing. For example, interventions focusing on retraining learned body-level OP mappings or compensating for specific disconnections within the reading network could be tailored to individual lesion profiles. Future research is needed to explore how lesion-induced network disruptions influence dynamic recovery processes to further refine neurorehabilitation strategies.

## Acknowledgments

This work is supported by the National Institute on Deafness and other Communication Disorders (NIDCD grants R01DC014960 and R01DC020446 to PET, K99DC018828 to ATD, F30DC018215 to JVD, and T32DC019481 to SMD). We thank the valuable contributions of our participants, and our data collectors, in alphabetical order: Elizabeth Dvorak, Trini Kelly, Elizabeth Lacey, Alycia Laks, Sachi Paul, and Candace van der Stelt.

## Supplemental Materials

**Supplemental Table 1:**
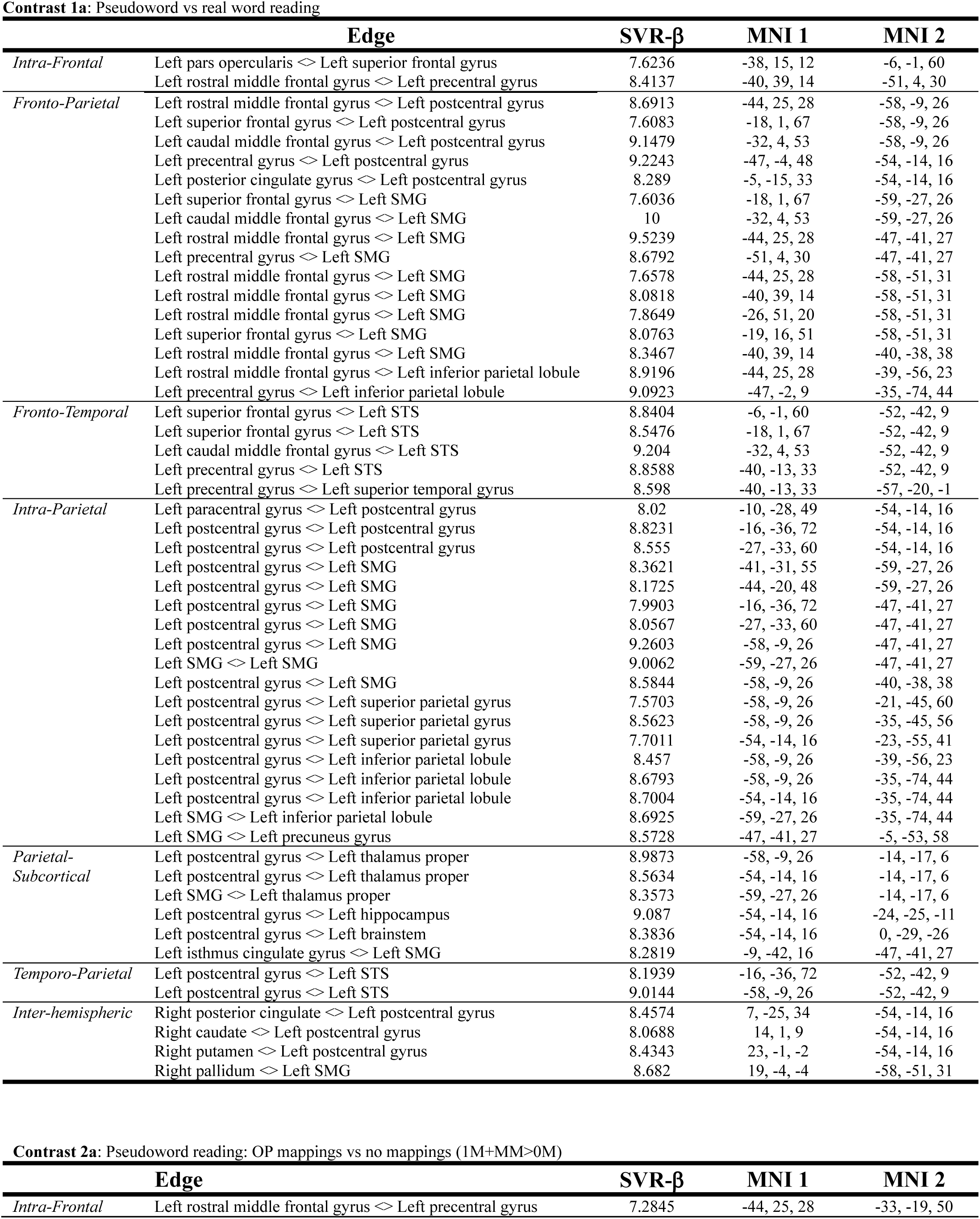
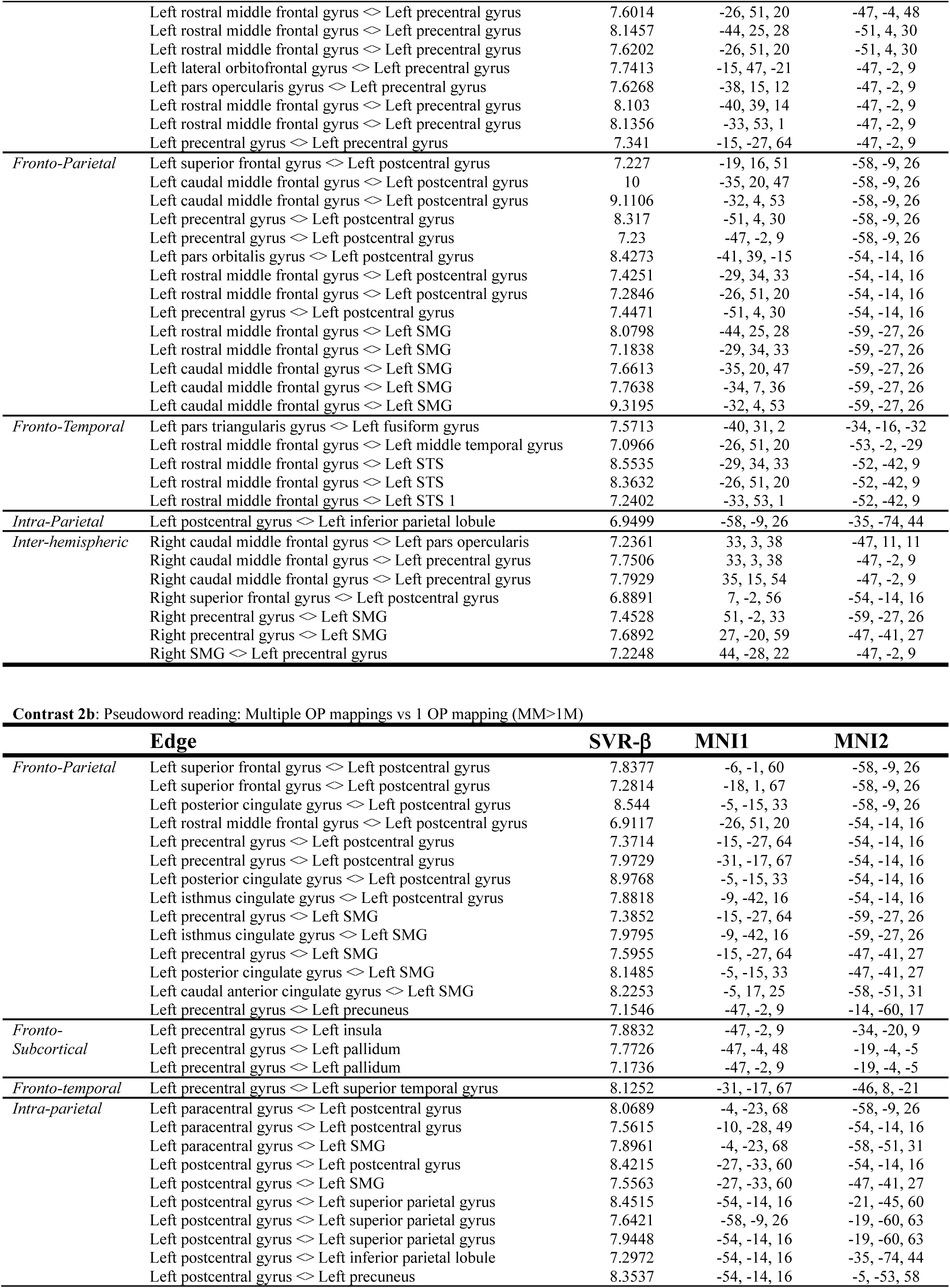
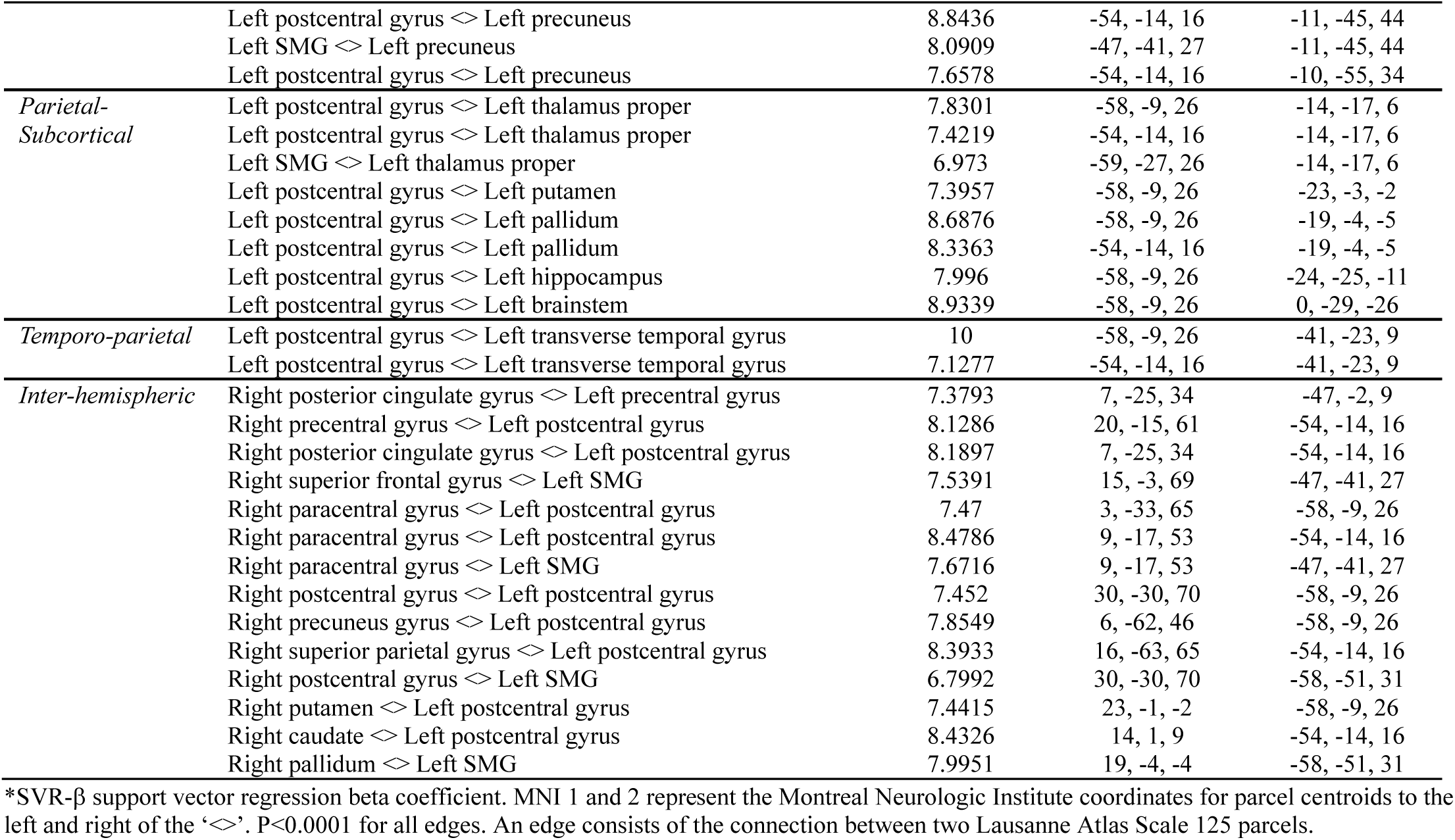
SVR-CLSM results for oral pseudoword reading accuracy in contrasts 1a, 2a, and 2b.

**Supplemental Table 2:**
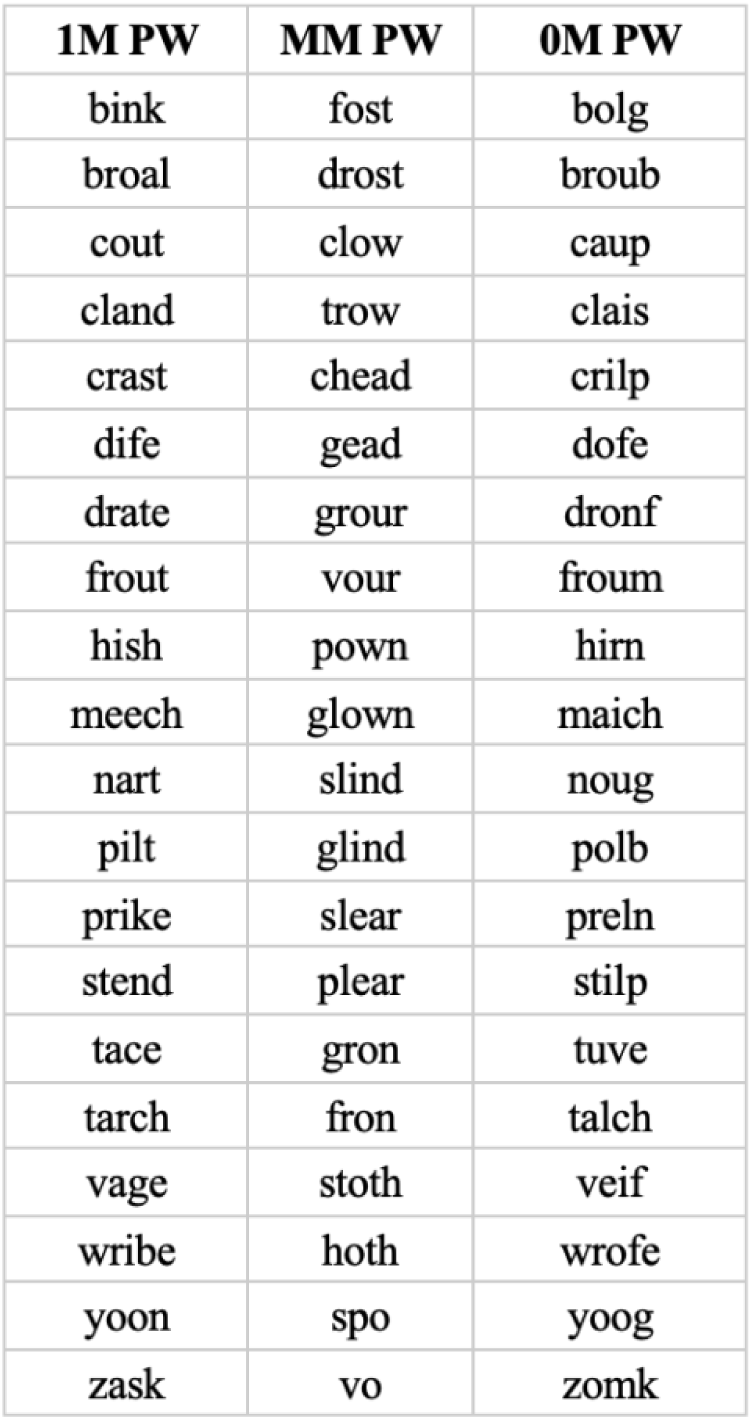
Oral Pseudoword (PW) Reading Stimuli by Type.

